# Prenatal Arsenic Exposure and Gene Expression in Fetal Liver, Heart, Lung, and Placenta

**DOI:** 10.1101/2024.11.10.622821

**Authors:** K. A. Rychlik, C. Kashiwagi, J. Liao, A. Mathur, E. J. Illingworth, S. S. Sanchez, A. Kleensang, A. Maertens, F. C. M. Sillé

**Author notes:** (corresponding author; address: Johns Hopkins Bloomberg School of Public Health, 615 N. Wolfe st. RM E7628, Baltimore, MD 21205;).

## Abstract

Prenatal arsenic exposure has been linked to a myriad of negative health effects. There is relatively little insight into the mechanisms and signaling alterations across different fetal organs that drive long-term immune-related issues following prenatal arsenic exposure. Therefore, the effects of this exposure window on gene expression in the liver, placenta, heart, and lung of gestation day (GD) 18 C57BL/6 mouse fetuses were investigated. From two weeks prior to mating until tissue collection at GD18, mice were exposed to 0 or 100 ppb sodium (meta) arsenite in drinking water. Genes of interest were analyzed by RT-qPCR, complemented with untargeted Agilent 44K microarray analysis. Data cleanup and analysis was performed in RStudio. Differentially expressed mRNAs were queried in the String Database and using Cytoscape to create interaction networks and identify significantly enriched biological pathways. A total of 251, 165, 158, and 41 genes were significantly altered in the liver, placenta, heart, and lung, respectively, when treated samples were compared to controls. Many altered pathways were immune-related, supporting prior research. Most notably, gene expression of Gbp3, a key player in the cellular response to interferon gamma, was found to be reduced in placentas of female fetuses exposed to arsenic compared to controls (*p*=0.0762).

**Impact:** This is the first study comparing alterations in gene expression across multiple organs following prenatal exposure to environmentally relevant levels of arsenic. These findings, elucidating the multi-organ impact of prenatal arsenic exposure on predominantly immune-related pathways, further our mechanistic understanding of the long-term health effects observed in early-life arsenic-exposed populations.

## Introduction

Despite strong population-level data supporting the association between prenatal inorganic arsenic exposure and cancer of the liver and lung, heart disease, and immune dysfunction (Smith et al. 2012; Naujokas et al. 2013; Khan et al. 2020), the mechanisms driving these long-term effects are still unclear. Arsenic is the most common chemical drinking water contaminant worldwide and puts roughly 230 million people at risk (Shaji et al. 2021), including the most vulnerable populations of pregnant women and children. Within the U.S. alone, an estimated 2.1 million people are drinking water contaminated with levels of arsenic higher than the recommended limit of 10 ppb (Ayotte et al. 2017).

In an effort to isolate the mechanisms driving effects, some work has turned to investigating the placenta. Identified as a route of early exposure to arsenic (García Salcedo et al. 2022), the placenta is a mediator of *in utero* exposure and, in the case of heavy metals, could provide an indication of altered signaling in the neonate due to maternal exposure (Myllynen et al. 2005). Early evidence, albeit at rather high doses (20 ppm), indicate disruption of placental vasculogenesis (He et al. 2007) which could result in placental insufficiency, a condition which, like arsenic, has been associated with fetal (or intrauterine) growth restriction (IUGR) (Neerhof and Thaete 2008; Xu et al. 2022).

In a U.S. cohort study of low-to-moderate levels of arsenic exposure during pregnancy, alterations in gene expression and the epigenetic profile of placental tissue were discovered with differences noted between male and female fetal sex (Winterbottom, Ban, et al. 2019; Winterbottom, Moroishi, et al. 2019). Although not identified in that cohort, one pathway that has been presented as a possible mediator for IUGR after prenatal arsenic exposure is the glucocorticoid receptor signaling pathway (Caldwell et al. 2015). Not only is this pathway essential for many endocrine functions but it is also plays a key role in metabolism and immune function (Kadmiel and Cidlowski 2013). Dose-dependent effects on glucocorticoid signaling has also been identified in JEG-3 cells, an *in vitro* model of trophoblasts (Meakin et al. 2019). Specifically, Meakin et al. (2019) identified DNA methylation as a driver of effects within that pathway. Epigenetic alterations and gene expression changes have also been identified in newborn mouse liver tissue following *in utero* exposure to 85 ppm arsenic (Xie et al. 2007).

In human cohorts, gene expression and cytokine levels have been affected in cord blood based on maternal arsenic exposure levels (Fry et al. 2007; Rager et al. 2014). In an integrative study of data from twelve prior cohort studies in humans looking at CpG methylation, gene expression, or protein expression in the placenta or cord blood, Laine & Fry (2016) determined that many of the commonly altered genes were related to immune and inflammatory pathways.

The question of whether gene expression alterations in the placenta at birth correspond to alterations in other organs and whether those alterations are mechanistic drivers for long-term disease, however, has yet to be answered. Therefore, this research set out to determine what gene expression alterations, if any, occurred following preconception and prenatal exposure to inorganic arsenic in drinking water among placenta, liver, heart, and lung tissues in the C57Bl/6J mouse model. Initially, genes of interest were selected to quantify using RT-PCR in each organ based on prior research and networks of interest, particularly in the intersection between the immune system and glucocorticoid signaling. Secondarily, a microarray was performed on a subset of samples to investigate larger network alterations and trends.

## Materials and Methods

### Animal Care and Mouse Exposure Model

Despite some interspecies differences, C57BL/6 mouse models have been proven representative for assessing arsenic immunotoxicity (Ramsey et al., 2013a; Ramsey et al., 2013b; Kozul et al., 2009). Breeder male and female C57Bl/6J mice (8 weeks of age) were obtained from Jackson Laboratories (Bar Harbor, Maine). Mice were given a week to acclimate prior to experiments. Male mice were housed individually. Female mice were housed individually at the onset of arsenic exposure in order to accurately monitor food and water intake. Enrichment was added to cages including a mouse igloo and additional sterile paper bedding material. Cages were housed in a temperature and humidity-controlled facility with 14:10 light:dark cycle and provided *ad libitum* access to food and water. Low arsenic chow diet was obtained from Research Diets Inc. (AIN-93M). ***Exposure model:*** Mice were evenly distributed (by weight) into two groups two weeks prior to mating: Control with access to plain water (0 ppb iAs) and Exposure with access to exposure water (spiked with 100 ppb iAs - a relevant dose in many areas of the world ( Ayotte et al. 2017; García Salcedo et al. 2022; Naujokas et al. 2013) - in the form of sodium(meta)arsenite (CAS# 7784-46-5) obtained from Millipore Sigma [St. Louis, MO]). All water was provided to the mice in glass containers to avoid risk of phytoestrogen interference in the study. Crystal Springs brand water (Lakeland, FL), with reported non-detectable levels of arsenic, was refreshed every 3-4 days to avoid arsenic oxidation. No significant differences in water or food consumption were noted between exposure groups. Exposure paradigm is represented in **Figure 1**. ***Breeding strategy:*** To obtain fetuses, timed matings were carried out by exchanging female and male bedding three days prior to mating, and then pairing 1:1 male:female in the male cage overnight. The next morning was designated GD0.5 (gestational day 0.5) and males were removed to avoid asynchronous pregnancies. Successful pregnancy was confirmed by weight gain. Fetuses were harvested from the first litter of pregnant females euthanized by CO_2_ overdose on gestational day (GD) 18. Placenta, lung, liver, and heart tissue were collected from fetuses and flash frozen using liquid nitrogen. ***Sample size:*** The total number of experimental conditions yielded 24 unique samples: 2 different exposure groups (unexposed vs. 100 ppb inorganic arsenic exposed) * 4 different tissues (fetal heart, liver, lung and placenta) * 3 animals per exposure group. Based on prior literature in other comparable studies (Torres, et al., 2016; Vilmundarson et al., 2022), biological triplicates (N=3) for each tissue per exposure group was determined to be sufficient for the current analysis. Each biological replicate represented a unique litter. ***Randomization:*** fetuses and corresponding placenta were randomly chosen from each litter (n=3 per exposure group) for RNA extraction. Sex as a biological variable was not determined nor considered during the randomization process. ***Inclusion & exclusion criteria:*** Only pregnant females were included in this study to obtain access to fetal tissues. Tissue samples would have been excluded from analysis if contamination occurred or if RNA yield would be less than the 100 ng required for the microarray, or if RNA quality was poor (pure was considered 260/280 ratio >2.0; 2.0 <260/230 ratio< 2.2; and 260/230 ratio > 260/280 ratio). No animals, experimental units, or data points were excluded for this analysis. ***ARRIVE Statement:*** This study was conducted with approval by the Johns Hopkins University Institutional Animal Care and Use Committee (Protocol # MO20H283), following the National Research Council’s Guide for the Care and Use of Laboratory Animals. In addition, we have been following the ARRIVE guidelines 2.0 (Percie du Sert 2020) for the entire study.

**Figure 1.**
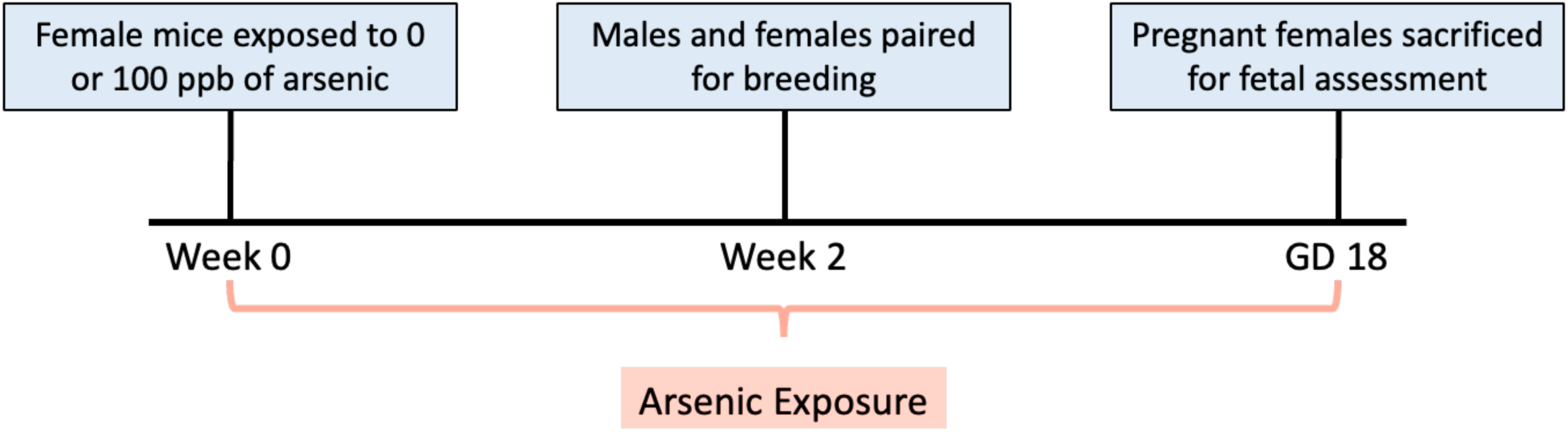
Preconception and prenatal arsenic exposure paradigm. C57Bl/6J mice were exposed to either 0 ppb (control) or 100 ppb (exposed) sodium (meta) arsenite (iAs) in drinking water from two weeks prior to breeding until euthanasia of the pregnant dam at gestational day (GD) 18. Samples were collected at GD 18 including liver, heart, lung, and placenta and flash frozen using liquid nitrogen.

### Total RNA Isolation

Tissues were thawed in RNAprotect (Qiagen Inc., Valencia, CA), ground using frosted slides and run through a Qiashredder column (Qiagen Inc., Valencia, CA) before RNA isolation. Total RNA was extracted from each organ (<30 mg) using RNeasy Mini kits (Qiagen Inc., Valencia, CA). Quantity and purity of total RNA was measured using a Take3 Microvolume Plate and the Take3 app (Agilent Technologies, Santa Clara, CA). Total RNA was considered pure when 260/280 ratio >2.0; 2.0 <260/230 ratio< 2.2; and 260/230 ratio > 260/280 ratio.

### RT-qPCR

RT-qPCR was run on a set of samples (N=2-5) using Universal SYBR Green Fast qPCR Mix (Cat no. RK21203; ABClonal Inc., Woburn, MA) and primer sets in the published literature (**Table 1**). All runs were normalized to the 18S ribosomal housekeeping gene (**Table 1**) previously used by Espindola et al. (2012). For proteasome 20S subunit beta 8 (PSMB8), a TaqMan probe was utilized (Cat no. 4453320; Applied Biosystems, Waltham, MA) following instructions from the manufacturer.

**Table 1.**
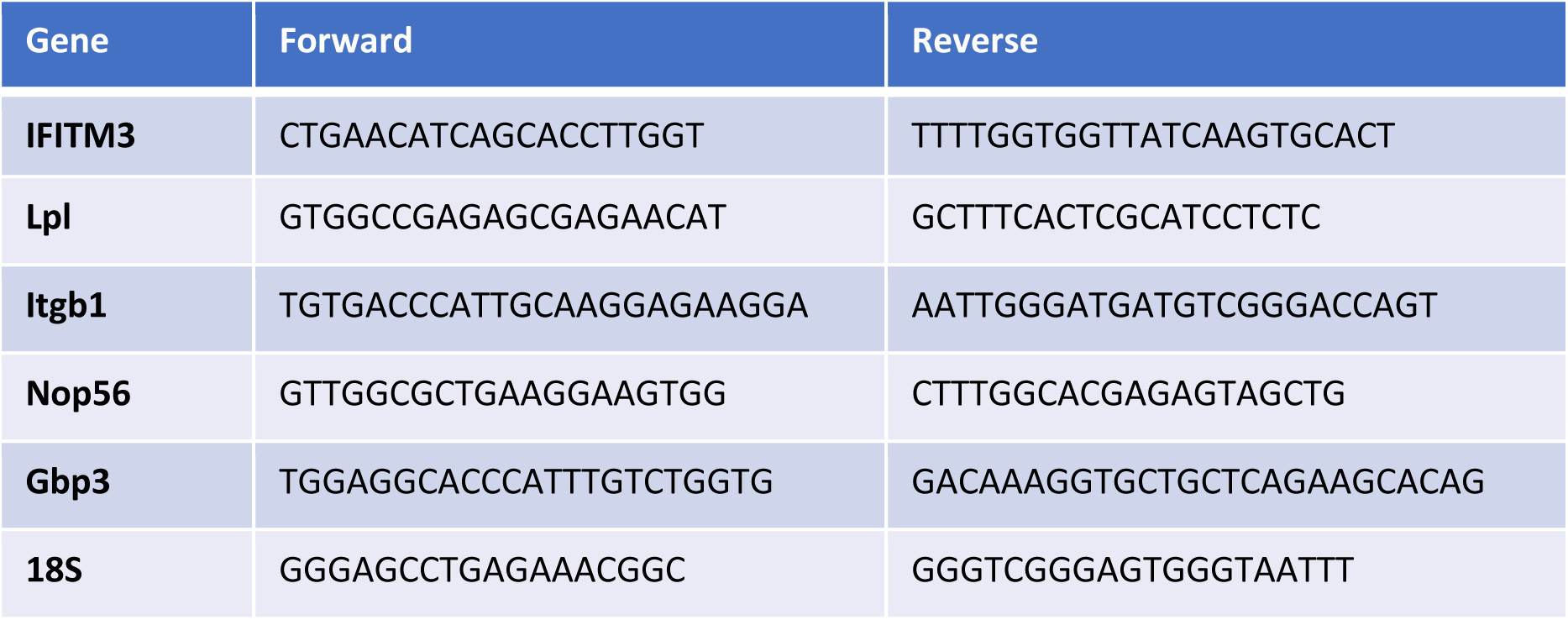
Primer sets for RT-qPCR. The primer sets listed in the table were utilized for analysis of mRNA for individual genes selected from the most dysregulated genes from the microarray data that demonstrated the most connections in generated STRING maps. All mRNA quantification was normalized to 18S ribosomal mRNA expression.

### Microarray

Total RNA was reverse-transcribed to cDNA using the Applied Biosystems High-Capacity cDNA Reverse Transcription Kit (Cat no. 4368814; Applied Biosystems, Waltham, MA).

Gene expression analysis was carried out using Agilent Whole Mouse 44K Genome Oligo Array (Agilent Technologies) according to manufacturer instructions. Due to poor RNA isolation, the total number of samples included in the final analysis was N=3 per group (arsenic & control).

### Statistical Analysis

RT-qPCR results were analyzed using GraphPad Prism 9 (GraphPad Software, Boston, MA). Groups were compared using an ANOVA with post hoc analysis Bartlett’s statistic (corrected; for Guanylate-Binding Protein 3) or Tukey’s multiple comparisons test (for all others).

Data from microarray experiments were preprocessed using GeneSpring (Agilent Technologies, Santa Clara, CA) software. Raw data were imported and quantile-normalized. Statistical analysis was done using limma with a moderated Bayes t-test and a Benjamini-Hochberg correction. Although no individual gene was statistically significantly changed after correcting for multiple hypothesis testing, all genes with an unadjusted *p*-value less than 0.01 were investigated for further pathway analysis.

### Protein-protein Interaction Networks and Pathway Enrichment

We queried over-represented pathways using Search Tool for the Retrieval of Interacting Genes/Proteins (STRING, v12.0, string-db.org), a database that extracts experimental and curated data from several sources to deduce protein-protein interaction (PPI) networks from a set of protein-coding genes (Szklarczyk et al. 2023). Using a differentially expressed genes (DEGs) list cutoff, |log2FC|>0 and adjusted *p-*value cutoff of <0.05, we generated PPI networks for the gene sets for each tissue using the STRING medium confidence required interaction score. When no individual gene was statistically significantly changed after correcting for multiple hypothesis testing, all genes with an uncorrected p-value less than .01 were investigated for over-represented pathways via STRING. These were visualized in STRING with all potential interaction sources used and a moderate level of evidence; enrichment statistics were performed using STRING with the assumption of the whole genome as background. We limited our network generation to only include experimental evidence and evidence from curated databases as active interaction sources. Experimental databases in STRING are (BIND, DIP, GRID, HPRD, IntAct, MINT, and PID), and the curated databases in STRING include the gene ontology (GO) database (Gene Ontology Consortium 2004), KEGG database (Kanehisa and Goto, 2000), Biocarta, BioCyc and Reactome. Finally, Cytoscape (v 3.9.1) (Shannon et al. 2003) an open-source platform, was used to search whether there were any additional pathways.

## Results

### qRT-PCR Analysis

Selected genes based on prior arsenic exposure literature were analyzed using RT-PCR to examine organ-specific alterations in gene expression following preconception and prenatal arsenic exposure. Although no significance was discovered, there was a trending association in placenta. In the liver, IFITM3 and PSMB8 were unchanged when treatments were compared to controls (**Figure 2, A & B**). Unfortunately, the number of control liver samples were limited in this sample set; therefore, any alterations that may exist could have been clouded by a small sample size. Similarly, in the heart, no significant differences were observed in relative expression of lipoprotein lipase (Lpl) or integrin beta 1 (Itgb1) (**Figure 2, C & D**). One sample in the male arsenic group had higher expression of 18S ribosomal RNA compared to other samples (still within two standard deviations of the mean) which resulted in much higher 2^ΔΔCt values for both heart mRNAs. Nop56 ribonucleoprotein (Nop56) expression in lung tissue was also unaffected by treatment, with some samples showing higher expression than others, but no major trends were observed across treatment groups (**Figure 2E**). Finally, Guanylate-Binding Protein 3 (Gbp3) expression in placenta neared significance when comparing female arsenic-exposed placentas to controls (**Figure 2F**). Arsenic-exposed samples exhibited higher levels of this mRNA (*p*=0.0762).

**Figure 2.**
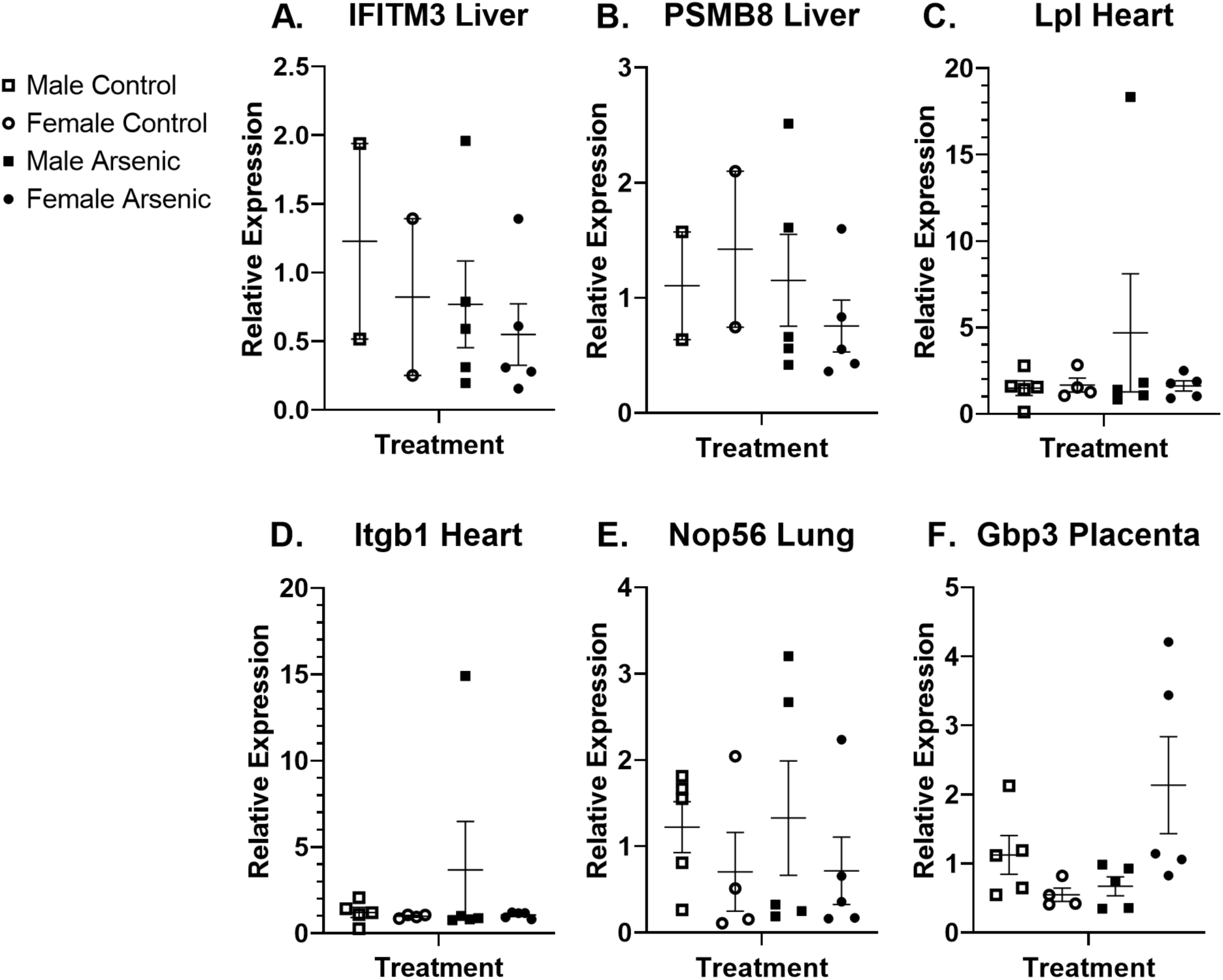
RT-qPCR Analysis of Select Genes in Lung, Liver, Heart, and Placenta of GD18 mice offspring prenatally exposed to arsenic. Tissues including heart, lung, liver, and placenta were excised upon euthanasia of the pregnant dam at gestational day (GD) 18 following preconception and prenatal exposure to either 0 ppb (control) or 100 ppb (exposed) sodium (meta) arsenite. Tissues from the C57Bl/6J mice were flash frozen using liquid nitrogen and stored at -80° C. RNA was extracted using Qiagen kits, reverse transcribed, and gene expression analysis was carried out using RT-qPCR. Select genes (IFITM3 and PSMB8 in liver, Lpl and Itgb1 in heart, Nop56 in lung, and Gbp3 in placenta) were analyzed by RT-qPCR to examine organ-specific alterations in gene expression. All runs were normalized to 18S ribosomal RNA. N=2-5 from at least two litters. No findings reached statistical significance.

### Microarray Analysis

Raw data from microarray experiments was preprocessed, quantile-normalized and differential gene expression between exposed (100 ppb iAs) and unexposed (0 ppb) was determined for fetal heart (**Table S1**), liver (**Table S2**), lung (**Table S3**) and placenta (**Table S4**). Of the genes altered with an unadjusted *p*-value <0.01, just four mapped genes were changed across all tissues: the heart, liver, and placenta (Pim3, Ifit3, Psmb8, and Rtp4; **Figure 3**). Between the heart and liver, overlap occurred in 15 genes: C1qb, Tyrobp, Hic1, Ankrd37, Ly6c1, Gm16430, Isg15, Ifitm3, Gcm1, Irf9, Usp18, Oas1a, Oasl1, Xaf1, and Gzma. Heart and placenta had eight of the same differentially expressed genes (Hmga1, Irgm2, Irgm1, Tmem140, Pde4b, Ifi204, Gbp3, and Cmpk2). Liver and placenta had just three overlapping genes (Tgtp2, Pfkfb3, and Dek). While 49 genes were found to be differentially expressed in the lung, none overlapped with any of the other organs studied. After the altered genes for each organ were input into STRING and a map was acquired, dysregulated biological pathways were identified.

**Figure 3.**
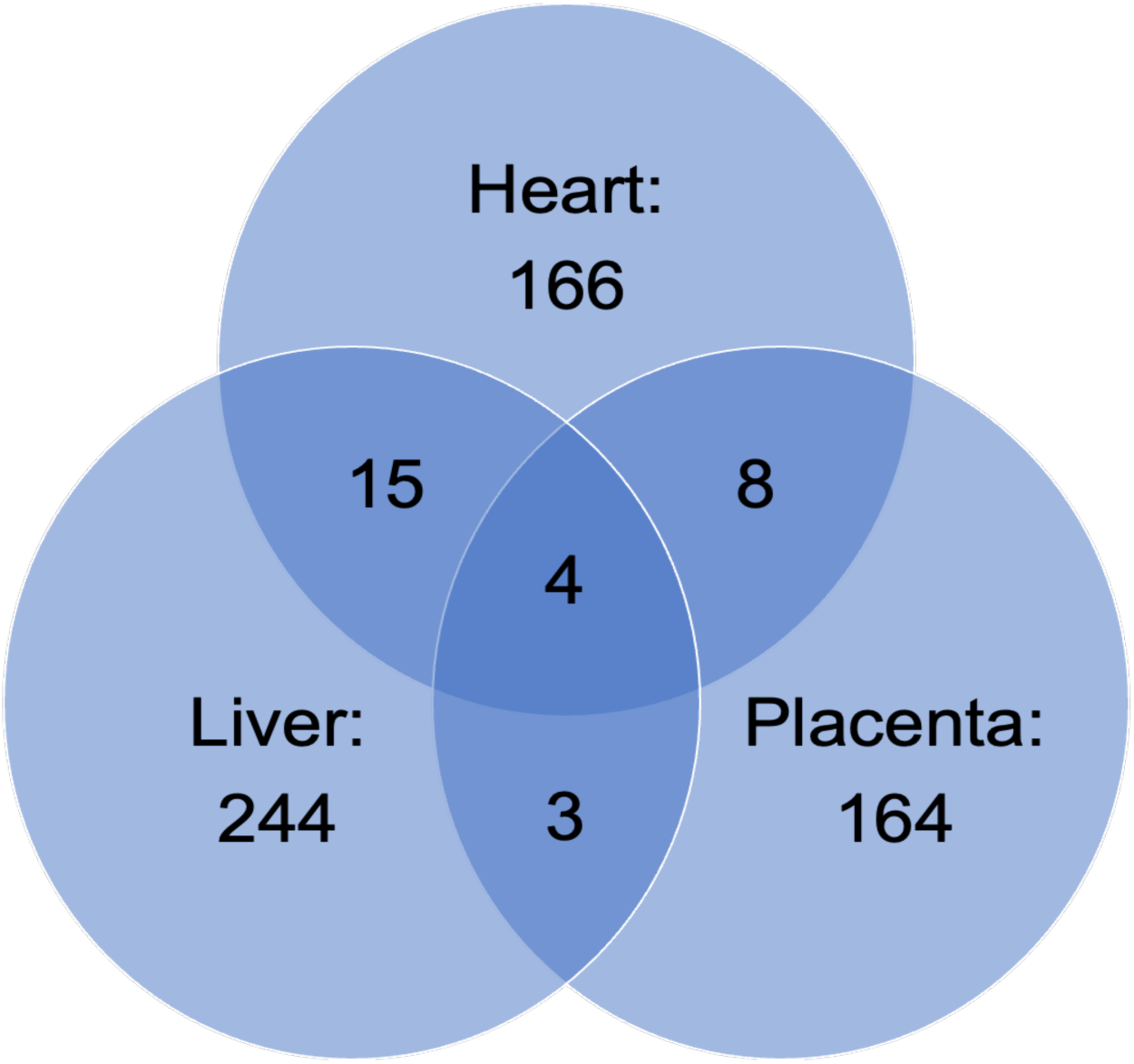
Differential gene expression overlap between heart, liver, and placenta. Tissues including heart, lung, liver, and placenta were excised upon euthanasia of the pregnant dam at gestational day (GD) 18 following preconception and prenatal exposure to either 0 ppb (control) or 100 ppb (exposed) sodium (meta) arsenite. Tissues from the C57Bl/6J mice were flash frozen using liquid nitrogen and stored at -80° C. RNA was extracted using kits from Qiagen, reverse transcribed, and gene expression analysis was carried out using the Agilent Whole Mouse 44K Genome Oligo Array. Among the genes altered with an unadjusted *p*-value <0.01, only four were found to overlap between placenta, liver, and heart. Similarly, there was minimal overlap between heart and liver (15), liver and placenta (3), and heart and placenta (8). No overlapping genes were identified between the lung and any other organ. Data shown represents an N=3.

#### Fetal Liver

Findings from the liver revealed 251 differentially expressed transcripts, of which 240 were known and could be input into STRING (**Figure 4**). In the microarray, Interferon-induced transmembrane protein 3 (Ifitm3) was downregulated in arsenic-exposed mice (-1.02 log fold change). Interestingly, Ifitm3 and Psmb8 were two of the genes found to be highly connected to other altered transcripts. Enriched biological processes included regulation of biological process, cellular process, and biological regulation (top 10 results in **Table 2**, complete list in **Table S5**). No KEGG pathway enrichment was identified in the STRING database based on the input. After inputting the same differentially expressed transcripts into Cytoscape, the human pathway related to immune response to tuberculosis (WP4197) was identified. In particular, the downstream regulation of this pathway includes PSMB8, IFIT3, and interferon regulatory factors (IRF) 1 & 9, which were found to be differentially regulated in the current mouse model.

**Figure 4.**
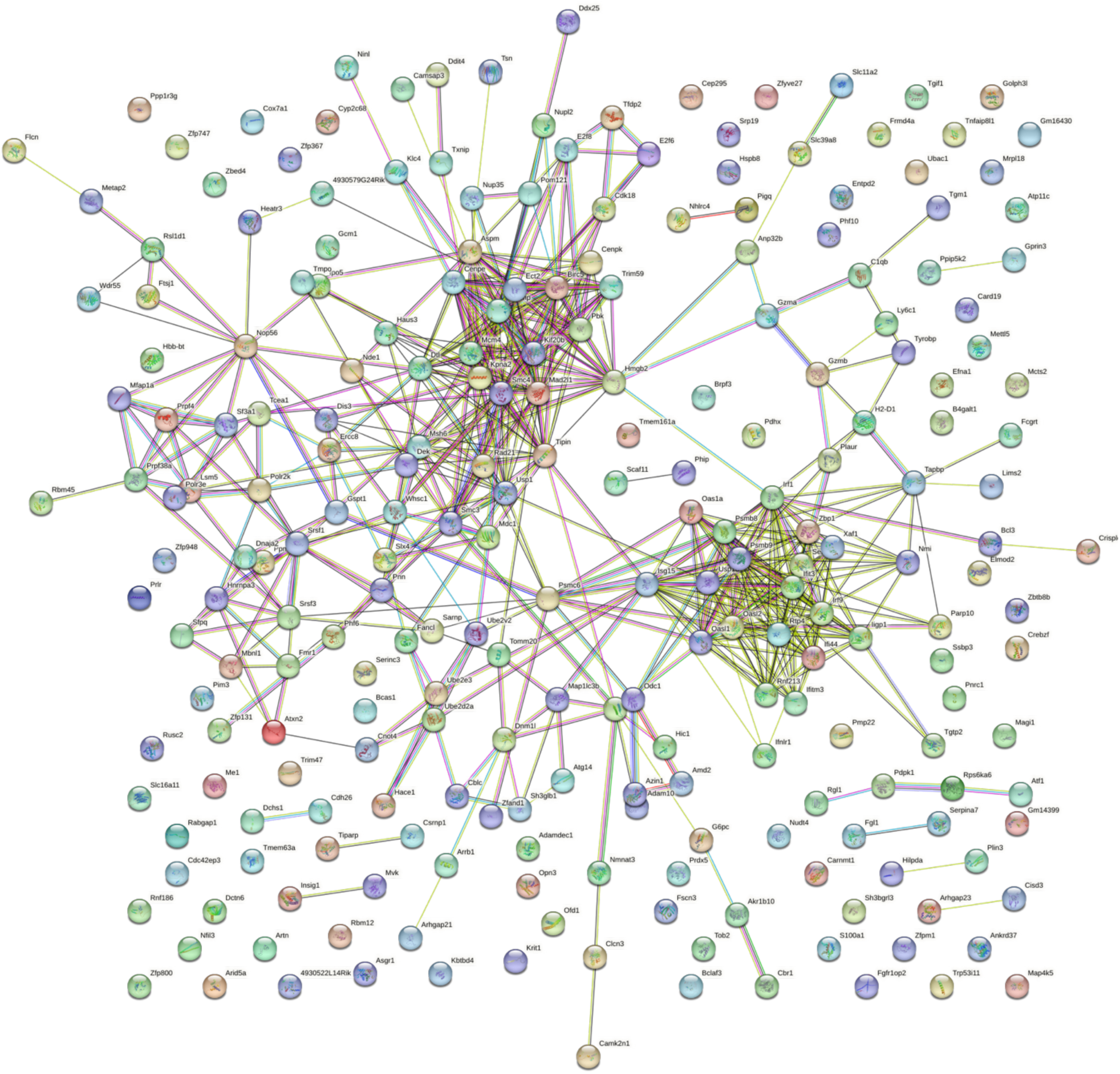
Liver STRING Map. Liver samples were obtained from gestational day (GD) 18 C57Bl/6J fetal mice exposed for 2 weeks prior to conception and during gestation to either 0 ppb (control) or 100 ppb (exposed) sodium (meta) arsenite. After microarray analysis, 240 differentially expressed (comparing exposed to controls exhibited an unadjusted *p*-value <0.01) genes from the liver were entered into the STRING database to generate a map. Nodes are indicated by the colored circles; lines indicate connections between nodes. Data shown represents an N=3.

**Table 2.**
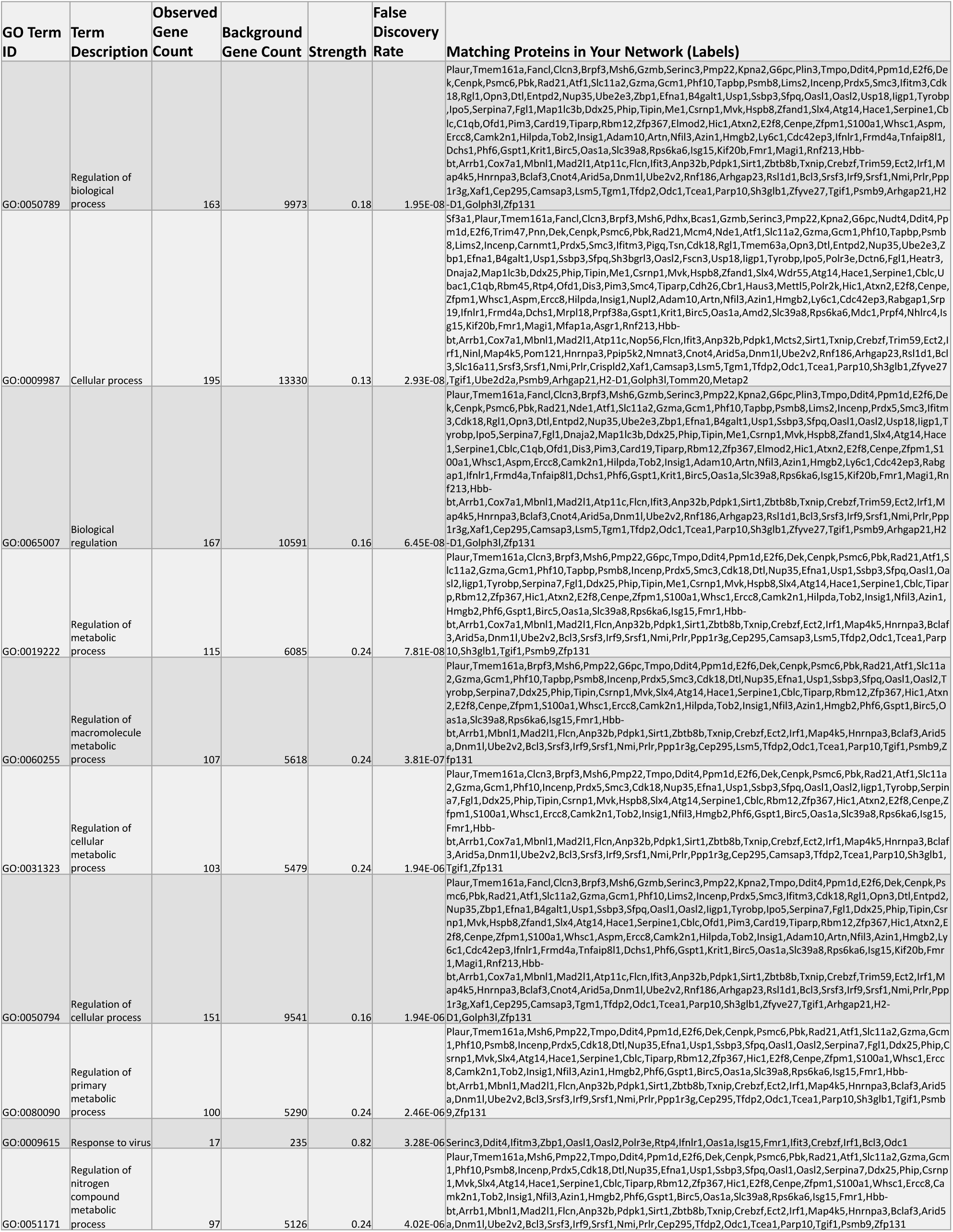
Top 10 Enriched Biological Processes in arsenic-exposed GD18 Liver Tissue. Along with generation of the STRING map, enriched biological processes were identified based on the 240 differentially expressed (comparing exposed to controls exhibited an unadjusted *p*-value <0.01) genes from the liver using gene ontology (GO). The specific GO number is reported along with its term description. Each protein from the differentially expressed input that are associated with that biological process are listed in the right column of the table. Data shown represents an N=3.

#### Fetal Heart

Data from the heart revealed 163 differentially expressed RNA transcripts, of which 150 known transcripts were entered into STRING. A STRING interaction map was generated (**Figure 5**) and information was obtained on enriched biological processes (based on gene ontology -GO) and KEGG pathways (top 10 STRING/GO findings in **Table 3**, all KEGG findings in **Table 4**). The three most enriched biological processes were immune system process, immune response, and immune effector process. Similarly, enriched KEGG pathways revolved mainly around immune response with the top three as prion diseases, osteoclast differentiation, and the B cell receptor signaling pathway. Again, in the heart, Psmb8 was a highly connected transcript, as was Ifitm3. C-C motif chemokine ligand 5 (Ccl5) appeared to be a key factor linking various components of the two major networks identified in the STRING diagram. Both Itgb1 and Lpl were slightly outside of the main networks but still connected to many other dysregulated genes. Cytoscape inquiry revealed the human macrophage markers CD68 and coagulation factor 3 (F3) as two of the dysregulated genes related to the WikiPathway WP4146 – Macrophage markers.

**Figure 5.**
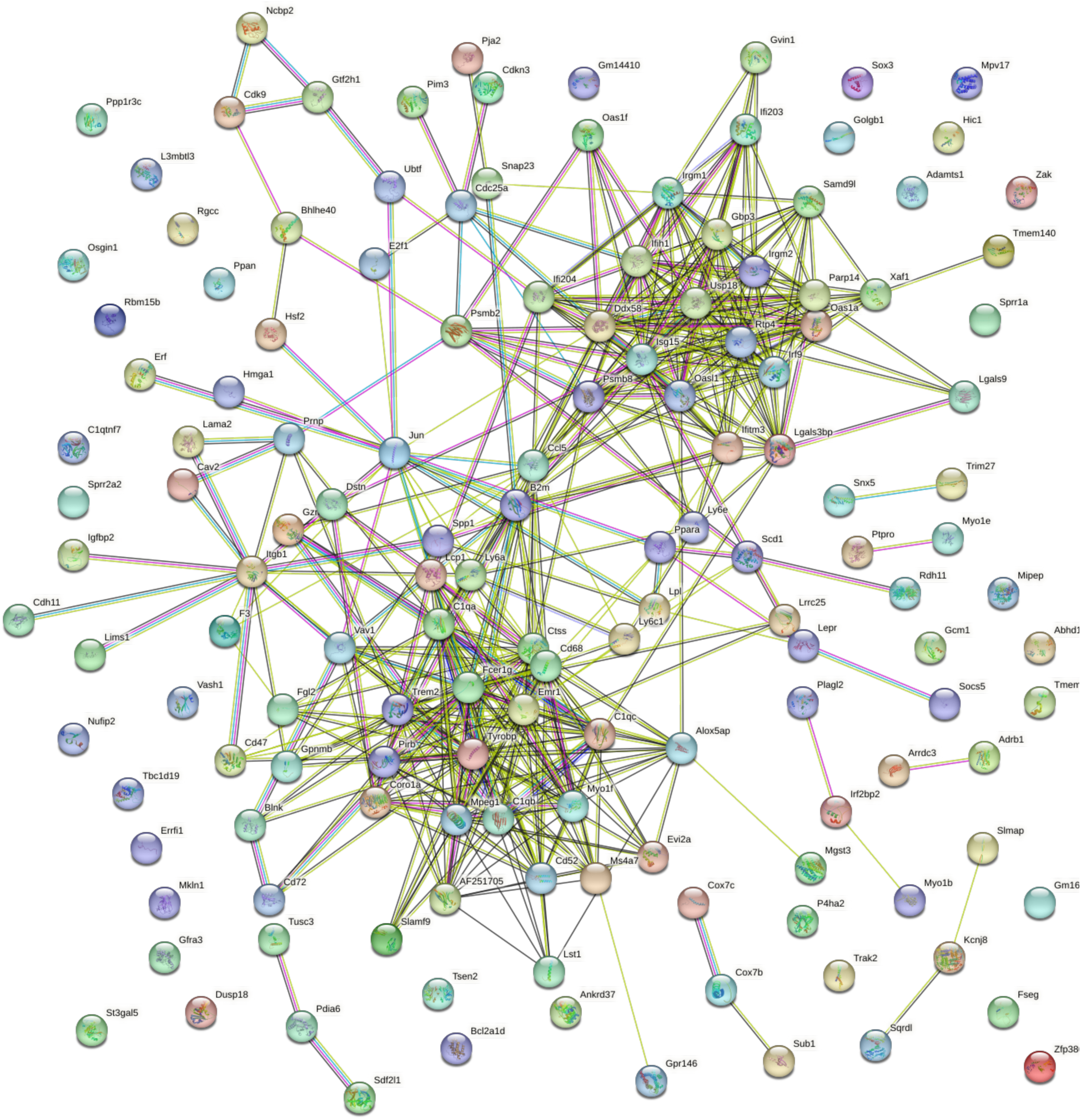
Heart STRING Map. Heart samples were obtained from gestational day (GD) 18 C57Bl/6J fetal mice exposed for 2 weeks prior to conception and during gestation to either 0 ppb (control) or 100 ppb (exposed) sodium (meta) arsenite. After microarray analysis, 156 differentially expressed (comparing exposed to controls exhibited an unadjusted *p*-value <0.01) genes from the heart were entered into the STRING database to generate a map. Nodes are indicated by the colored circles; lines indicate connections between nodes. Data shown represents an N=3.

**Table 3.**
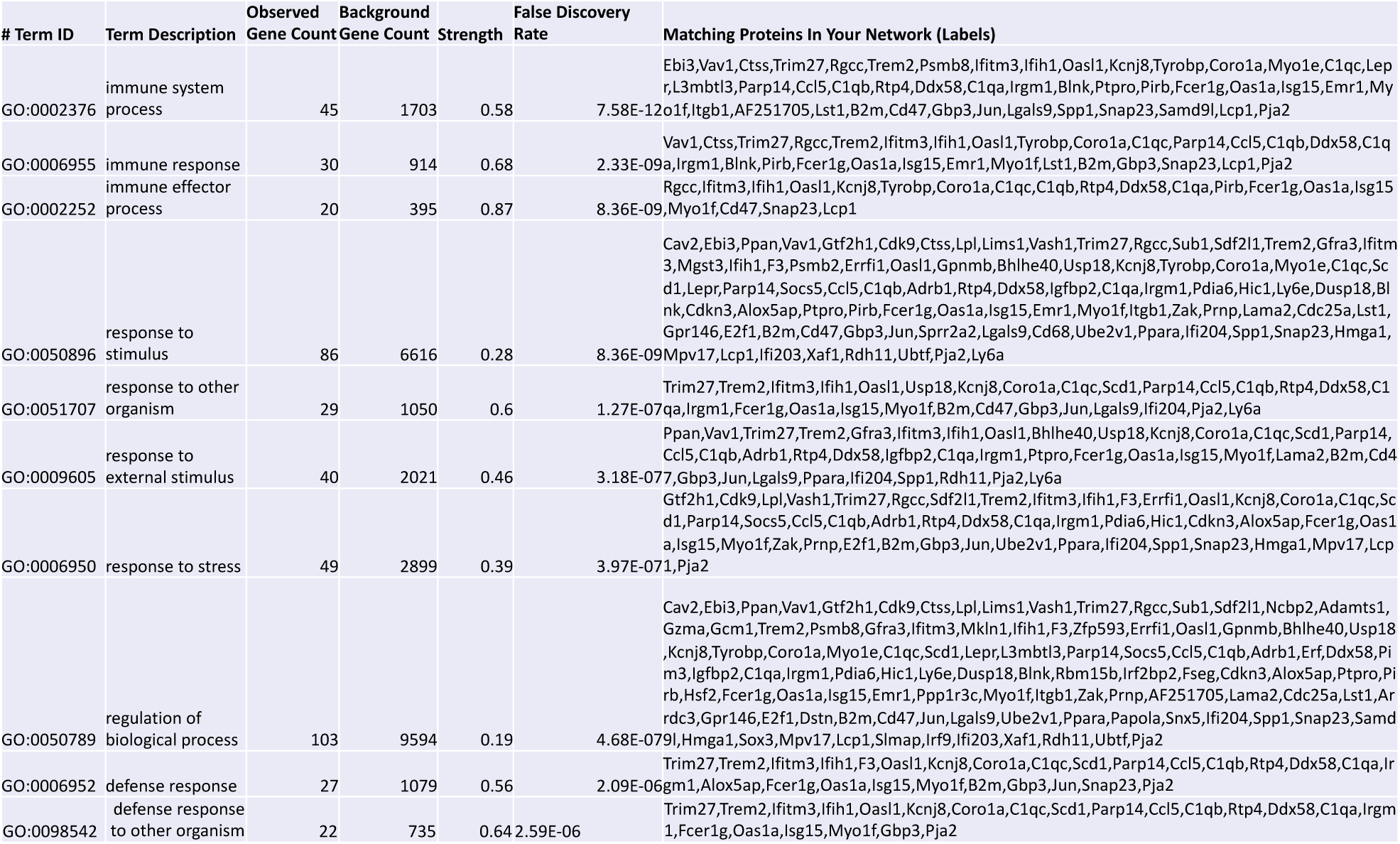
Top 10 Enriched Biological Processes in arsenic-exposed GD18 Heart Tissue. Along with generation of the STRING map, enriched biological processes were identified based on the 150 differentially expressed (comparing exposed to controls exhibited an unadjusted *p*-value <0.01) genes from the heart using gene ontology (GO). The specific GO number is reported along with its term description. Each protein from the differentially expressed input that are associated with that biological process are listed in the right column of the table. Data shown represents an N=3.

**Table 4.**
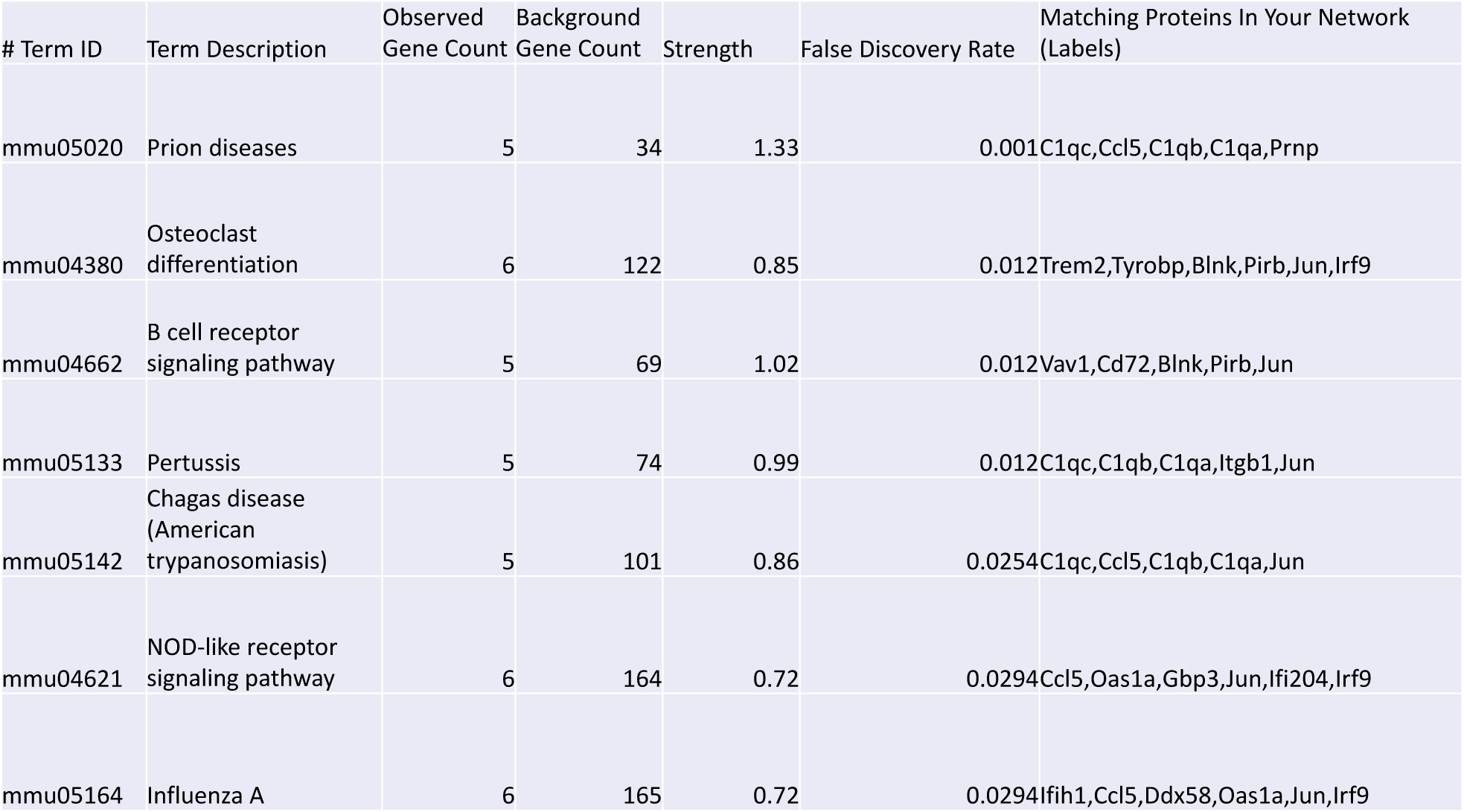
Enriched KEGG Pathways in arsenic-exposed GD18 Heart Tissue. Enriched KEGG pathways were also identified by STRING based on the 150 differentially expressed (comparing exposed to controls exhibited an unadjusted *p*-value <0.01) genes from the heart. The specific term identification number is reported along with its term description. Each protein from the differentially expressed input that are associated with that pathway are listed in the right column of the table. Data shown represents an N=3.

#### Fetal Lung

A total of 49 mRNAs were altered in lung tissue, with 41 of these being input into the STRING program. Nop56 was one of the most connected transcripts identified in the generated STRING map for the lung. Although a small network of genes was identified to be interacting (**Figure 6**), four biological processes were found to be enriched within the set including chromatin assembly, ribosome biogenesis, cellular component biogenesis, and nucleosome assembly (**Table 5A**). Just one KEGG pathway was enriched; cysteine and methionine metabolism (**Table 5B**). Similarly, analysis in Cytoscape mapped DNA methyltransferase 3B (Dnmt3b) and 3-phosphoglycerate dehydrogenase (Phgdh) to WP2525, the pathway for trans-sulfuration and one-carbon metabolism.

**Figure 6.**
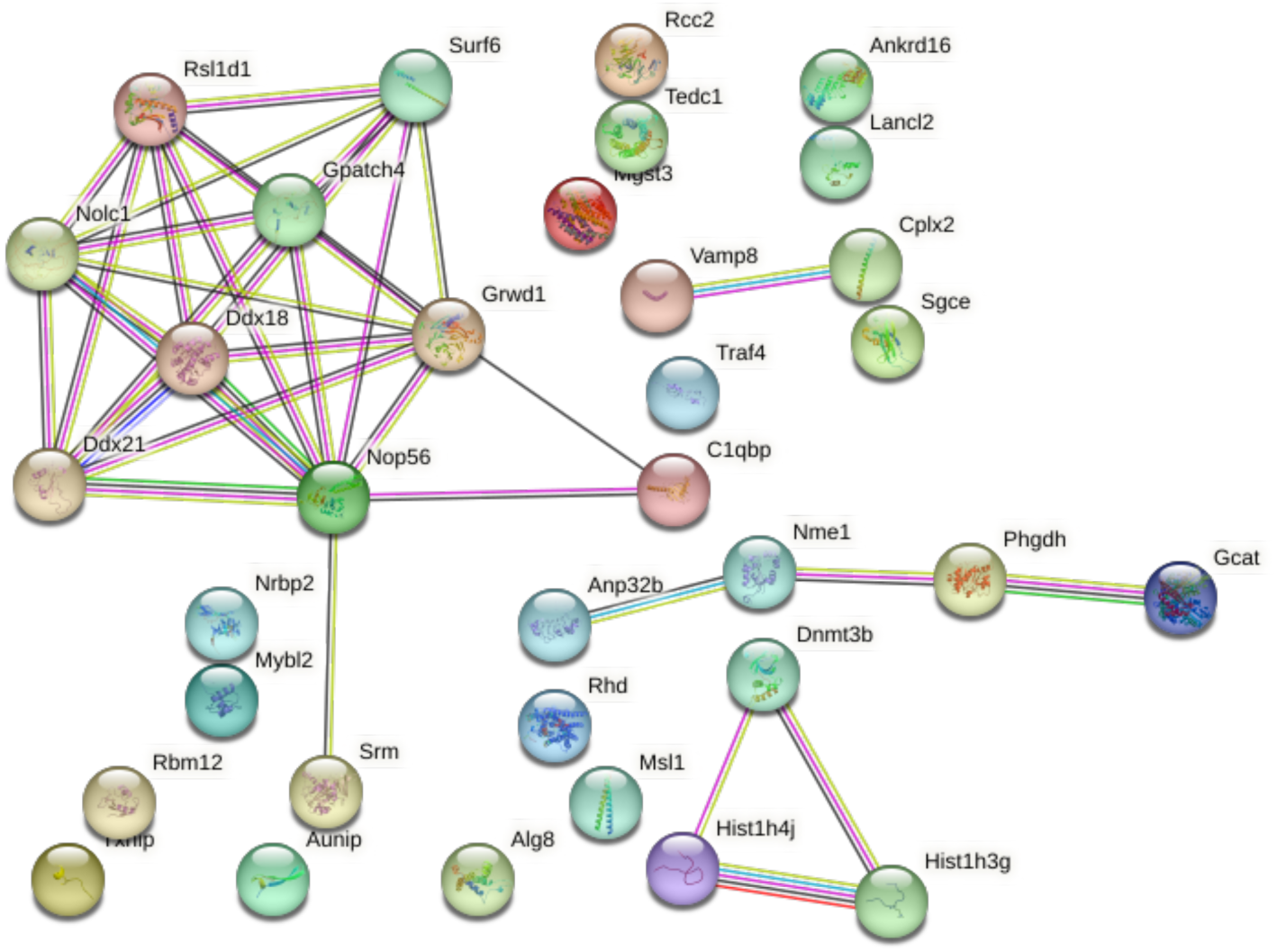
Lung STRING Map. Lung samples were obtained from gestational day (GD) 18 C57Bl/6J fetal mice exposed for 2 weeks prior to conception and during gestation to either 0 ppb (control) or 100 ppb (exposed) sodium (meta) arsenite. After microarray analysis, 41 differentially expressed (comparing exposed to controls exhibited an unadjusted *p*-value <0.01) genes from the lung were entered into the STRING database to generate a map. Nodes are indicated by colored circles; lines denote connections between nodes. Data shown represents an N=3.

**Table 5.**
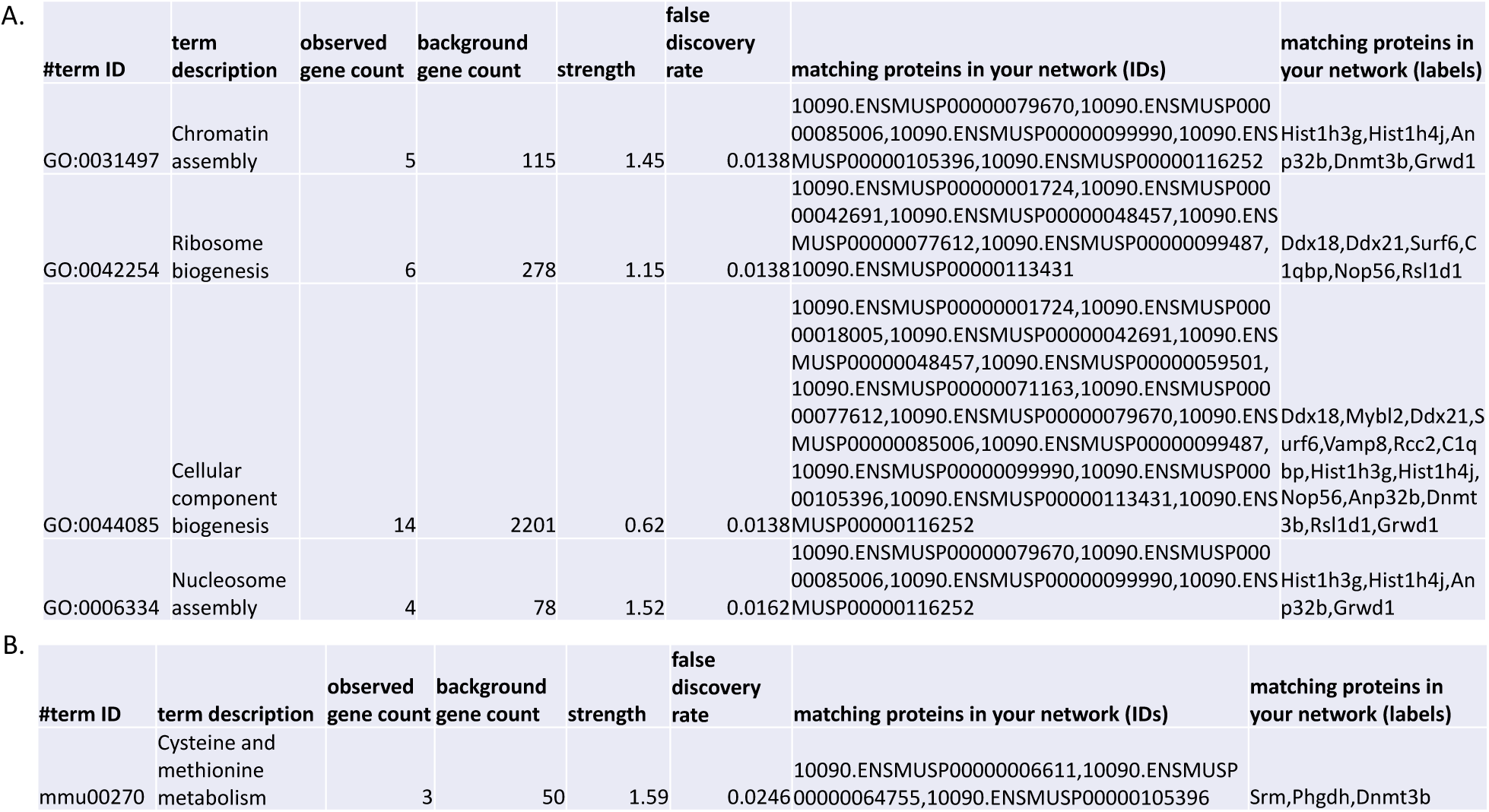
Enriched Biological Processes (A) and Enriched KEGG Pathways (B) in arsenic-exposed GD18 Lung Tissue. Along with generation of the STRING map, **(A)** enriched biological processes and **(B)** KEGG pathways were identified based on the 41 differentially expressed (comparing exposed to controls exhibited an unadjusted *p*-value <0.01) genes from the lung using gene ontology (GO) and KEGG terms. The specific GO number or KEGG term ID number is reported along with its term description. Each protein from the differentially expressed input that are associated with that biological process or pathway are listed in the right column of the tables. Data shown represents an N=3.

#### Placenta

In the placenta, there were 190 differentially expressed mRNAs; 165 of these were input into STRING because many of them were not mapped to a known transcript. There was one more highly connected area in the STRING map and many of those connections included Gbp3 and Psmb8 (**Figure 7**). Although there were no GO or KEGG pathways found to be enriched in this data, enriched cellular components were identified (**Table 6**). After input into Cytoscape, the same pathway identified for liver tissue exhibited similarity in the dysregulated genes for placenta tissue. Specifically, Ifit3, Psmb8, and protein tyrosine phosphatase, non-receptor type 2 (Ptpn2) mapped to this pathway.

**Figure 7.**
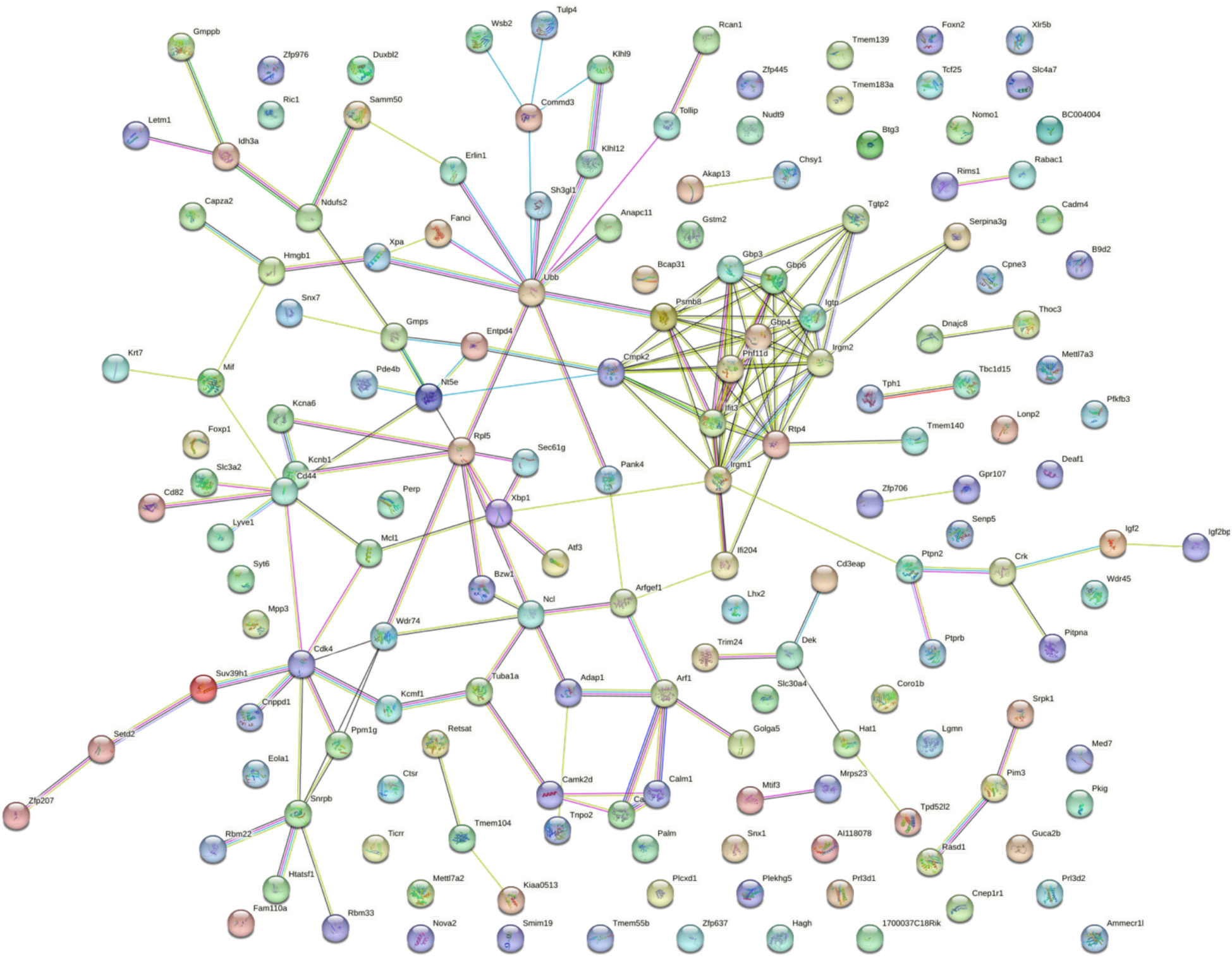
Placenta STRING Map. Placenta samples were obtained from gestational day (GD) 18 C57Bl/6J fetal mice exposed for 2 weeks prior to conception and during gestation to either 0 ppb (control) or 100 ppb (exposed) sodium (meta) arsenite. After microarray analysis, 165 differentially expressed (comparing exposed to controls exhibited an unadjusted *p*-value <0.01) genes from the placenta were entered into the STRING database to generate a map. Nodes are indicated by colored circles; lines denote connections between nodes. Data shown represents an N=3.

**Table 6.**
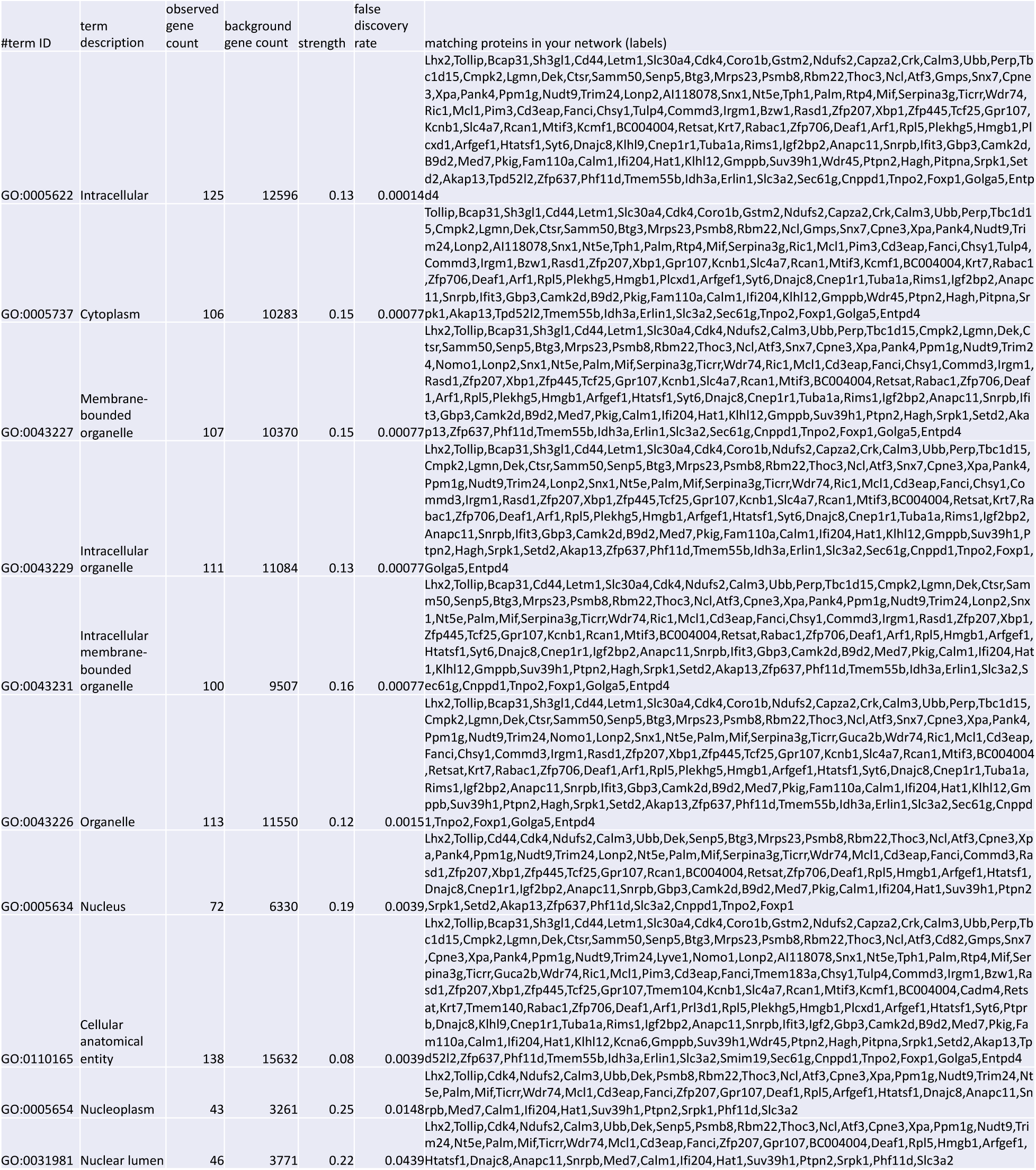
Top 10 Enriched cellular components in arsenic-exposed GD18 Placenta Tissue. Along with generation of the STRING map, enriched cellular components were identified based on the 165 differentially expressed (comparing exposed to controls exhibited an unadjusted *p*-value <0.01) genes from the placenta using gene ontology (GO). The specific GO number is reported along with its term description. Each protein from the differentially expressed input that are associated with that biological process are listed in the right column of the table. Data shown represents an N=3.

## Discussion

Although Ifitm3 was not found to be significantly different in the liver of this model upon examination by RT-PCR, it was one of the most differentially expressed transcripts in the microarray and was highly connected in the STRING map. Considering that the microarray only included one sample, different to those run with RT-PCR, perhaps that individual was a high expresser for this transcript. However, it reveals that investigation of a single transcript in response to low level arsenic exposure is not sufficient to display a complete portrait of effects. Furthermore, in both the liver and placenta, this dysregulated gene, along with Psmb8, mapped to the pathway involved in immune response to tuberculosis. Ifitm3 normally acts to inhibit viral entry into host cell cytoplasm (Li et al. 2013; Narayana et al. 2015); therefore, as a potential mechanism of immune suppression following prenatal arsenic exposure, it is of interest for further investigation. Ifitm3 was one of just a few genes found to be altered in the heart, liver, and placenta and has been identified as an intrinsic factor which can be upregulated by interferons (Jiang et al. 2018). It is also recognized as a limiting factor in influenza pathogenesis (Everitt et al. 2012) and seems to play a role in human metastasis prevention (Gómez-Herranz et al. 2019). Found to be downregulated in chronically iAs-treated hamsters (Hernández et al. 2011), Ifitm3 was identified as a potential mediator of hepatic carcinogenesis. Despite the fact that the exposure level was significantly higher in the hamsters (15 mg/L or 15,000 ppb), Ifitm3 and related pathways represent potential targets for mechanistic intervention. Furthermore, given the data on long-term susceptibility to tuberculosis following prenatal exposure to arsenic, Ifitm3 and Psmb8 may be mediators, along with other intrinsic factors, of those long-term effects.

Psmb8, or proteasome 20S subunit beta 8, is a key component of MHC class II and is expressed in many tissues in the body (PubChem). It is induced by interferon gamma and, interestingly, mutation in this subunit of the immunoproteasome has been linked to increased inflammation and lipodystrophy (Kitamura et al. 2011). Results from the current inquiry reveal a small downregulation of Psmb8 in the placenta, heart, and liver. Prior work found no effect of arsenic trioxide on expression of Psmb8 in NB4 acute promyelocytic leukemia cells; however, in the same study, a reduction in promoter activity of Psmb8 was noted (Yang et al. 2014).

Despite setting out to discover the relationship between prenatal arsenic exposure and the glucocorticoid receptor signaling pathway, the results highlight the known link between arsenic and impaired lipid metabolism (Renu et al. 2018), which is, of course, also linked to macrophage function. Nop56, found to be dysregulated in the lung of this model, has previously been found to be associated with metal exposure (including arsenic) and atherosclerosis (Riffo-Campos et al. 2018). Prenatal exposure has previously been linked to altered signaling in the liver, contributing to earlier onset of atherosclerosis in the apolipoprotein E-knockout mouse model (States et al. 2012). Lpl encodes a water-soluble enzyme important in lipoprotein processing, an essential macrophage function. It was not found to be different after RT-PCR analysis, but was identified as an altered transcript in the microarray. Interestingly, Lpl was found to be down-regulated following arsenic exposure due to its suppressive effect (arguably, toxicity) to macrophages (Song et al. 2019). Interestingly, recent work has shown that a drug inhibiting lipase activity can reverse these effects (Lou et al. 2022).

Other findings were less specific but many still are related to the immune response. For instance, Itgb1, involved in cell adhesion and recognition for multiple processes including immune response, did not show significant differences in RT-PCR analysis but exhibited altered expression in the microarray results. Further, this member of the arrestin family of proteins was found to be positively associated with arsenic exposure in two Bangladeshi cohorts (Demanelis et al. 2019). Gbp3 (-0.43 log FC), a transcript previously found to be upregulated in arsenic-exposed fetal C3H mouse liver (Liu et al. 2007), is one of a group of GTPases upregulated in response to interferons. Evidence indicates that this variant acts to moderate influenza virus, decreasing viral activity within the cell (Nordmann et al. 2012). In this study, a decrease in Gbp3 was observed in the placenta and liver, in alignment with findings of larger immune pathway downregulation and increased susceptibility to respiratory infection observed in human populations following prenatal arsenic exposure (Farzan et al. 2016; Laine and Fry 2016).

Importantly, these findings underline the idea that although small changes may be observed in individual genes across organs, these minute alterations across a network can lead to larger ramifications at the system level. This idea of network dynamics is influencing the field of medicine and allowing for use of artificial intelligence in order to make predictions about disease and treatment based on transcriptomic data (Theodoris et al. 2023). Using the presented data, a comparison across organs, in conjunction with other studies allows for correlation and furthers understanding of the mechanistic basis for long-term effects of preconception and prenatal arsenic exposure.

To our knowledge this is the first publication making a comparison of gene expression alterations in multiple organs following prenatal exposure to a relevant level of arsenic. Further investigation into these pathways as potential mechanistic targets to prevent long-term effects of arsenic exposure is needed.

## Supporting information

Rychlik_Arsenic_microarray_Supplemental_Data

## Acknowledgements

We acknowledge Tyrone Howard for his assistance with animal care and tissue collections. We acknowledge Vy Tran for help with the data analysis. K.A.R. was affiliated with the Department of Environmental Health and Engineering, Bloomberg School of Public Health, Johns Hopkins University at the time of the research and is currently affiliated with the School of Health Professions, University of Mary Hardin-Baylor.

## Supplemental Data

**Table S1. Complete List of Differential Gene Expression Analysis for Heart Tissue.** Heart samples were obtained from gestational day (GD) 18 C57Bl/6J fetal mice exposed for 2 weeks prior to conception and during gestation to either 0 ppb (control) or 100 ppb (exposed) sodium (meta) arsenite. After microarray analysis, the expression of all genes from heart tissue were compared for arsenic exposed to unexposed controls. Data shown represents an N=3.

**Table S2. Complete List of Differential Gene Expression Analysis for Liver Tissue.** Liver samples were obtained from gestational day (GD) 18 C57Bl/6J fetal mice exposed for 2 weeks prior to conception and during gestation to either 0 ppb (control) or 100 ppb (exposed) sodium (meta) arsenite. After microarray analysis, the expression of all genes from liver tissue were compared for arsenic exposed to unexposed controls. Data shown represents an N=3.

**Table S3. Complete List of Differential Gene Expression Analysis for Lung Tissue.** Lung samples were obtained from gestational day (GD) 18 C57Bl/6J fetal mice exposed for 2 weeks prior to conception and during gestation to either 0 ppb (control) or 100 ppb (exposed) sodium (meta) arsenite. After microarray analysis, the expression of all genes from lung tissue were compared for arsenic exposed to unexposed controls. Data shown represents an N=3.

**Table S4. Complete List of Differential Gene Expression Analysis for Placenta Tissue.** Placenta samples were obtained from gestational day (GD) 18 C57Bl/6J fetal mice exposed for 2 weeks prior to conception and during gestation to either 0 ppb (control) or 100 ppb (exposed) sodium (meta) arsenite. After microarray analysis, the expression of all genes from placenta tissue were compared for arsenic exposed to unexposed controls. Data shown represents an N=3.

**Table S5. Complete List of Enriched Biological Processes in arsenic-exposed GD18 Liver Tissue.** Enriched biological processes were identified in STRING based on the 240 differentially expressed (comparing exposed to controls exhibited an unadjusted *p*-value <0.01) genes from the liver using gene ontology (GO). The specific GO number is reported along with its term description. Each protein from the differentially expressed input that are associated with that biological process are listed in the right column of the table. Data shown represents an N=3.

